# Strong Selection is Necessary for Evolution of Blindness in Cave Dwellers

**DOI:** 10.1101/031872

**Authors:** Reed A. Cartwright, Rachel S. Schwartz, Alexandra L. Merry, Megan M. Howell

## Abstract

Blindness has evolved repeatedly in cave-dwelling organisms, and investigating the loss of sight in cave dwellers presents an opportunity to understand the operation of fundamental evolutionary processes, including drift, selection, mutation, and migration. The observation of blind organisms has prompted many hypotheses for their blindness, including both accumulation of neutral, loss-of-function mutations and adaptation to darkness. Here we model the evolution of blindness in caves. This model captures the interaction of three forces: (1) selection favoring alleles causing blindness, (2) immigration of sightedness alleles from a surface population, and (3) loss-of-function mutations creating blindness alleles. We investigated the dynamics of this model and determined selection-strength thresholds that result in blindness evolving in caves despite immigration of sightedness alleles from the surface. Our results indicate that strong selection is required for the evolution of blindness in cave-dwelling organisms, which is consistent with recent work suggesting a high metabolic cost of eye development.

## Introduction

Blindness has evolved repeatedly across taxa in caves, creating nearly a thousand cave-dwelling species and many more sub-populations (Culver et al., 2000; Dowling et al., 2002; Bradic et al., 2012; Coghill et al., 2014). Surprisingly, many populations of blind individuals experience some level of immigration, which would be expected to prevent the fixation of blindness in a newly established population (Avise and Selander, 1972; Bradic et al., 2012; Coghill et al., 2014). Thus, blind cave-dwelling populations of typically sighted species pose an interesting challenge to our understanding of evolutionary biology. Namely, how does significant population differentiation evolve despite homogenizing immigration?

Several hypotheses have been put forward to explain the evolution of blindness in cave-dwelling species. Darwin suggested that eyes would be lost by “disuse” (Darwin, 1859). We now consider this hypothesis the “neutral-mutation hypothesis” — random mutations can accumulate in genes or regulatory regions related to sight when, as in caves, there is no purifying selection to eliminate them. However, the accumulation of mutations causing blindness due to mutation pressure would take a longtime to result in fixation of blindness in populations on its own (Barr, 1968). Thus, genetic drift has been proposed to accelerate the evolution of blindness due to mutation pressure (Kimura and King, 1979; Borowsky, 2015; Wilkens, 1988). This hypothesis of relaxed selection appears to be supported by the observation of a high number of substitutions in putative eye genes in the blind forms of cavefishes (Hinaux et al., 2013; Protas et al., 2006; Gross et al., 2009). However, repeatedly developing blindness in cave populations simply by drift in isolation seems unlikely.

Relaxing selection that maintains the eye, however, also allows for other agents of selection to act on this trait (Lahti et al., 2009). The “adaptation hypothesis” suggests that there is a cost to an eye; thus, individuals without eyes have greater fitness when eyes are are not helpful, resulting in the eventual elimination of seeing individuals. This cost may either come from the energy required to develop a complex structure or due to the vulnerability of the eye (Barr, 1968; Strickler et al., 2007; Jeffery, 2005; Protas et al., 2007; Niven, 2008; Niven and Laughlin, 2008; Moran et al., 2015). Alternatively, positive selection may act on genes related to the eye if these genes act pleiotropically on traits that are beneficial in the dark. For example, in the Mexican tetra (*Astyanax mexicanus*) increased expression of Hedgehog (Hh) affects feeding structures, allowing better foraging in low light conditions (Jeffery, 2001,2005). Increased Hh signaling also inhibits pax6 expression, which results in eye loss during development (Yamamoto et al., 2004; Jeffery, 2005). Alternatively, cryptic variation may be maintained in normal conditions and expressed as blindness only in case of stress, such as entry into the cave (Rohner et al., 2013). When the cryptic variation is “unmasked”, it is then exposed to selection and could become fixed in the population.

Given that there is often gene flow from surface populations into caves, it seems that blind phenotypes should be lost unless selection for blindness is large (Avise and Selander, 1972). Recent work suggests a very high cost to developing neural tissue, including eyes (Moran et al., 2015). This cost, combined with pleiotropic effects, could lead to blindness despite immigration. However, the level of selection required to induce blindness in cave populations has not been quantified.

Here, we model the effects of migration, selection, and mutation to determine the conditions required for the evolution of blindness. This model allows us to explore migration-selection-mutation balance. Where previous theory have explored this balance more generally (Haldane, 1930; Wright, 1931,1969; Hedrick, 2011; Nagylaki, 1992; Yeaman and Otto, 2011; Yeaman and Whitlock, 2011; Bulmer, 1972), we address cavefish evolution specifically. The amount of selection required to oppose the force of immigration is high, but consistent with previous work on metabolic costs in novel environments and selection in other species. Interestingly, drift only impacts blindness in the cave population in a limited range of combinations of selection, dominance, and migration.

## Model and Analysis

### Assumptions

We consider two populations: surface-dwelling and cave-dwelling. We are interested in determining when the cave population will evolve blindness, i.e. become mostly comprised of blind individuals, as has occurred in numerous natural systems. We first assume that the surface and cave populations do not experience drift (i.e. populations are of infinite size). Additionally, immigration from the surface population into the cave affects the allele frequency in the cave, but emigration from the cave to the surface does not affect the surface population, as we assume that the surface population is significantly larger than the cave. Generations are discrete and non-overlapping, and mating is random. We track a single biallelic locus, where *B* is the seeing allele and where *b* is blindness allele. The frequency of *b* is denoted by *Q* ∈ [0,1] on the surface and *q* ∈ [0,1] in the cave. On the surface, we assume that blindness is strongly selected against, and *Q* is dictated by mutation-selection balance.

### Calculating the frequency of the blindness allele

Within the cave, the life cycle is as follows. (1) Embryos become juveniles and experience constant, directional selection with relative fitnesses of *w_bb_*= 1 + *s*, *w_Bb_*= 1 + *hs*, and *w_BB_* = 1, where *s* ≥ 0 and *h* ∈ [0,1]. (2) Juveniles migrate into and out of the cave such that a fraction *m* of adults come from the surface and 1 – *m* from the cave, where 0 ≤ *m* ≤ 1. (3) Adults generate gametes with one-way mutation, where 0 ≤ *u* ≤ 1 is the probability that a functional *B* allele becomes a non-functional *b* allele. (4) Gametes unite randomly to produce embryos. Given this life cycle, we calculate the allele frequency of the daughter generation (*q*′) via standard equations:

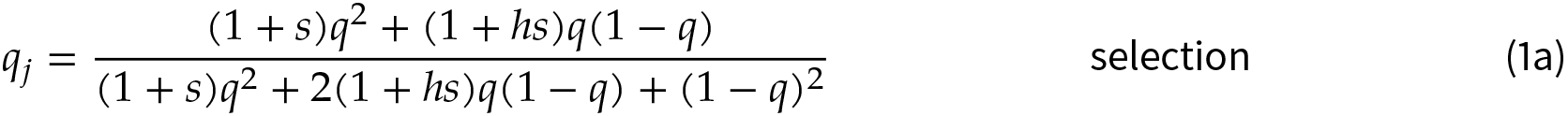

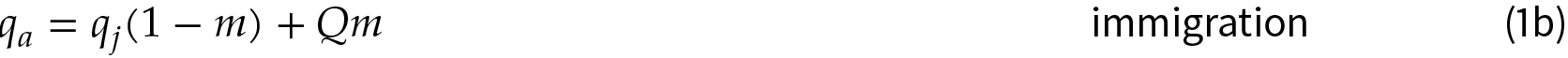

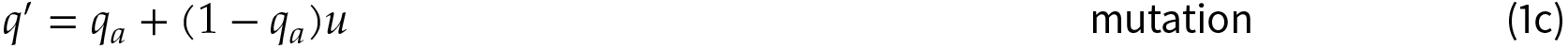

**Analysis of the change in allele frequency**. The change in allele frequency in one generation is Δ*q* = *q′ – q*, and each parameters influences it differently. Selection and mutation are directional forces, and increasing *s* or *u* increases Δ*q* for 0 ≤ *q* ≤ 1. (The derivatives are non-negative.) Increasing *h* causes selection to be more effective at low *q*, as rare *b* alleles are exposed to selection, but less effective at high *q*, as rare *B* alleles are sheltered from selection. Increasing *h* increases Δ*q* if 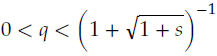 and decreases it if 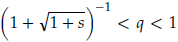. (The derivative is positive below this threshold and negative above it.) Migration harmonizes the allele frequency in the cave to the surface allele frequency. Thus increasing *m* increases Δ*q* for low *q* and decreases Δ*q* for high *q*. (The derivative is positive only when 0 ≤ *q* < *q*_*z*_(*h,s,Q*) ≤ *Q*, where *q*_*z*_ is a function describing a threshold.) However, increasing *Q* increases Δ*q* for 0 ≤ *q* ≤ 1. (The derivative is non-negative.)

### Identifying equilibriumallele frequencies

The model we have developed is an example of migration-selection balance (Wright, 1969; Hedrick, 2011; Nagylaki, 1992), extended to also include mutation. An equilibrium exists for this model when Δ*q* = 0. For small *s*, there is only one equilibrium, and it is near 1. For large *s*, there is only one equilibrium, and it is near 1. Three equilibria will only exist for moderate levels of selection (Figure 1). If *s* = *m* = *u* = 0, all forces of evolution are eliminated and Δ*q* = 0 for 0 ≤ *q* ≤ 1. A lower bound for any valid equilibrium is 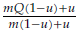 (Lemma 1). An upper bound for any equilibrium is 1 – *m*( 1 – *u*)( 1 – *Q*) (Lemma 2). Furthermore, it is important to note that

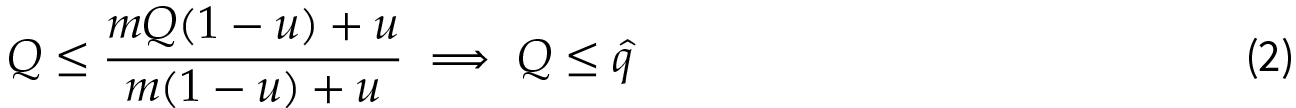

indicating that the equilibrium frequency in the cave will be greater than or equal to the allele frequency on the surface.

**Figure 1:**
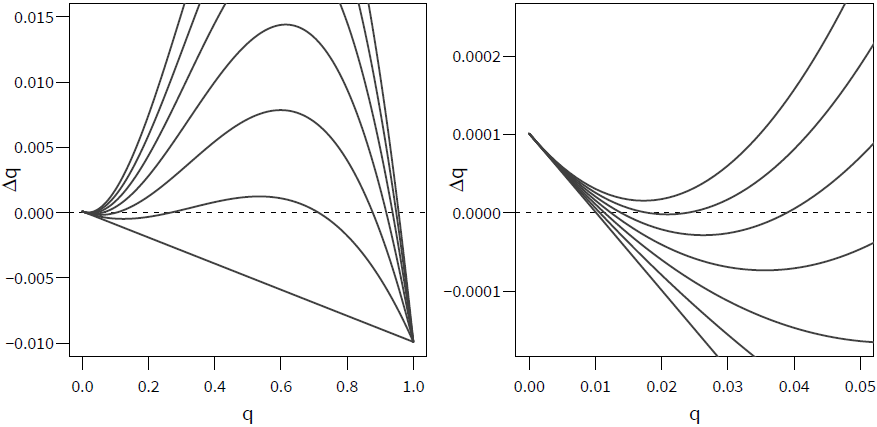
As selection increases, the evolutionary dynamics of the cave population changes. When *s* is low (bottom line; *s* = 0), there is only one equilibrium: near 0. As *s* increases (middle five lines, *s* = 0.05, 0.1, 0.15, 0.2, and 0.25) the local maximum (upper hump) increases and crosses the x-axis, producing three equilibria. When *s* gets high enough (top line; *s* = 0.3), the local minimum (lower valley) also crosses the x-axis, resulting in one equilibrium again. For all curves *m* = 0.01, *h* = 0, *u* = 10^-6^, and *Q* = 0.01. The figure on the right is an enlarged view of a small part of the figure on the left.

Assuming *s* > 0, the solution to Δ*q* = 0 are the roots of the following cubic polynomial

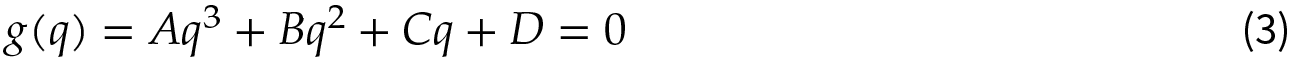

where

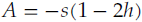

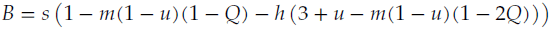

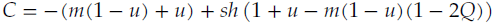

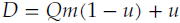

There are three possible roots of this equation, corresponding to three possible equilibria. Depending on the parameter values, Equation 3 may have three real roots or one real root and two imaginary roots. While the values of the roots of this polynomial can be expressed analytically, these equations are too complex to be helpful for understanding the system. For simplicity, we will let 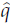 represent any possible equilibrium, and 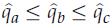 stand for the roots of Equation 3.

**Protected polymorphism**. Rather than tackling the equilibria directly, we first demonstrate that the cave has a protected polymorphism. A protected polymorphism exists if the allele frequency moves away from both fixation and extinction, i.e. Δ*q* > 0 when *q* = 0 and Δ*q* < 0 when *q* = 1. For *q* = 0, Δ*q* = *Qm*( 1 – *u*)+ *u* and *q* = 0 will be an equilibrium only if *Q*_*m*_ = 0 and *u* = 0; otherwise Δ*q* > 0 at *q* = 0. For*q* = 1, Δ*q* = −*m*(1 – *Q*)(1 – *u*) and *q* = 1 will be an equilibrium if *m* = 0, *Q* = 1, or *u* = 1; otherwise Δ*q* < 0. Thus a protected polymorphism always exists except at the edge cases *Qm* = *u* = 0, *m* = 0, *u* = 1, and *Q* = 1. In biological terms, the cave population will be polymorphic despite directional selection for *b* if there is some immigration from the surface population and the surface population is polymorphic.

**Validity of equilibria**. An equilibrium is only valid in our model if it is real and between [0,1]; otherwise, it is not biologically interpretable in this system. Because there is a protected polymorphism, there will be either 1 valid, stable equilibrium, or 3 valid equilibria in a stable-unstable-stable configuration, depending on the parameter values. While we have not exhaustively determined the parameter ranges under which there will be only one valid equilibrium, we have determined that if *h* ≥ 1/3 or if *h*< 1/3 and 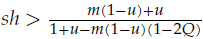, there will be only one valid equilibrium (Lemma 3).

We can also estimate the amount of selection required such that *q* is an equilibrium (*g*(*q*) = 0, Equation 3):

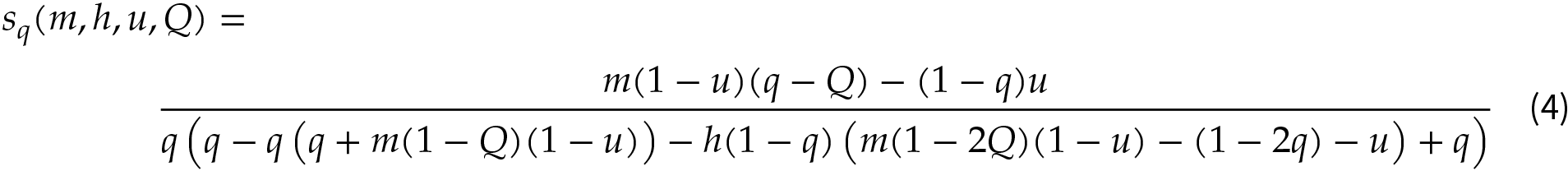

This equation is not valid for all *m* ∈ [0,1]. If the migration rate is low, 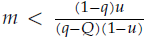, no level of selection will make *q* an equilibrium, as all equilibria will be greaterthan *q*. Similarly, if the migration rate is high,

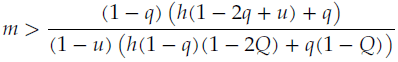

no level of selection will make *q* an equilibrium, as all equilibria will be less than *q*.

**Dynamics and the evolution of blindness**. The dynamics of the evolution of the cave population depend on the parameter values and the starting allele frequency, *q*_0_. If there is one equilibrium, then the frequency of *b* will evolve monotonically towards it, i.e. 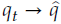 as *t* → ∞. If there are three equilibria, then the frequency of *b* will evolve monotonically to 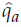 if 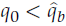 and to 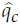 if 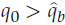.

When the cave population is founded, its initial allele frequency will likely match the equilibrium frequency on the surface (*q*_0_ = *Q*). Because 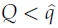 (Equation 2), the allele frequency in the cave will increase due to selection until it reaches the lowest equilibrium, i.e. *q_∞_*= inf{*q* : 0≤ *q* ≤ 1 and Δ*q* = 0}. Whether blindness evolves in the cave depends on whether *q_∞_* ≥ *q*\*, where *q*\* is a researcher-chosen threshold for determining that the cave population is a “blind” population. For example, *q*\* = 0.5 would specify that the blindness allele is the majority allele, and *q\** = 0.99 would determine that the blindness allele is approximately fixed. We can also focus on phenotypes, and let *a* = *q*^2^+2*q*(1 – *q*)*h* measure the average blind phenotype in the cave; then

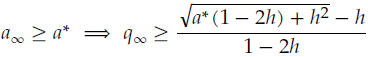

We define *s\** as the minimum level of selection required for cave population to become blind, given the other parameters, i.e.

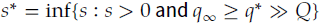

For simplicity, we will only consider values of *q*\* much higher than the surface allele frequency. If there is one equilibrium, 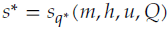; however, if there are three equilibria, *q*_*t*_ will evolve to the lower equilibrium and *q*_∞_ ≈ *Q* ≠ *q*\* (typically). Thus selection needs to be strong enough such that there is only one equilibrium; therefore,

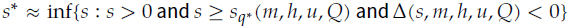

where Δ(*s,m,h,u,Q*) is the discriminant of Equation 3. Figure 2 plots analytical solutions for *s*\* based on different thresholds. When *m* ≫ *u*, the ratio *s*\*/*m* is roughly constant such that if *q_∞_* ≥ *q*\* then

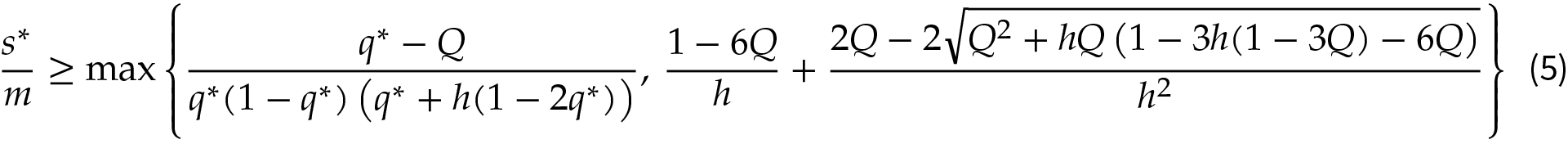

See Appendix for derivation.

**Figure 2:**
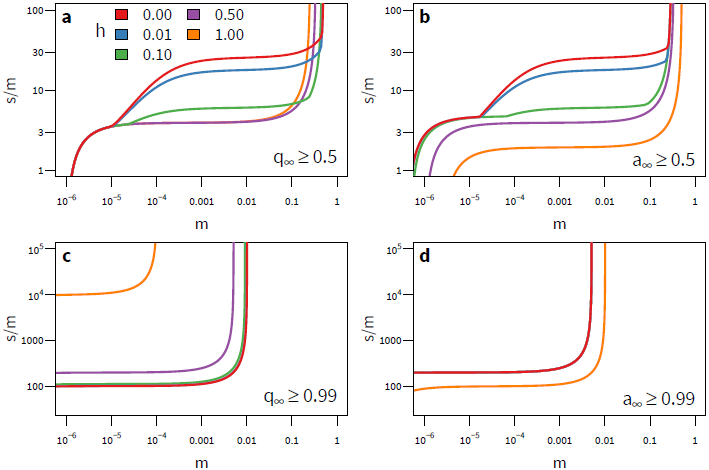
The level of dominance (*h*) of the blindness allele (*q*) affects the level of selection (*s*) required to produce blind populations. Each line represents how strong selection must be relative to migration (*m*) for blindness to evolve in the cave for a given level of dominance (*s*\*/*m*). Regions above the curves produce populations that are “blind” and regions below are not. Each panel contains a different condition for defining whether the cave is blind. (a) For the blind allele to become the majority allele requires stronger selection when the allele is recessive (*h* = 0). (b) For the blind phenotype to become the majority phenotype requires stronger selection when the allele is recessive. (c) For the blind allele to become fixed requires stronger selection when the allele is dominant. (d) For the blind phenotype to become fixed requires stronger selection when the allele is recessive. The curves were calculated analytically with *u* = 10.^-6^ and *Q* = 0.01.

### Recessive Blindness

In order to study the equilibria in more detail we limit subsequent work to a model where blindness is recessive (*h* = 0). As we have previously shown the effects of varying *h*, its impact on subsequent results can be inferred generally. First, we will simplify our model by assuming that *u* ≪ 1 such that 1 – *u* ≈ 1 and

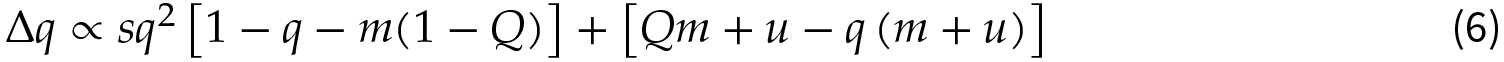

**Weak-selection approximation**. If selection is weak, then an equilibrium exists near *q =Q*. We use a second-order Taylor series at *q* =0 to determine the upper bound on *s* for the presence of three equilibria (i.e. when selection is so strong that an equilibrium near *Q* does not exist). The second-order series allows us to determine the lower two equilibrium points; although, this approximation is inaccurate as q increases. This approximation gives us

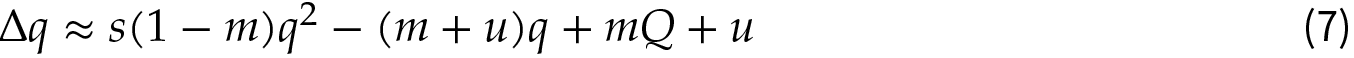

after assuming that 1 – *Q* ≈ 1. This equation has two roots, which are the lowest two of three total equilibria,

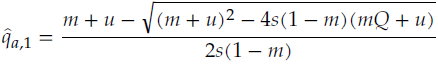

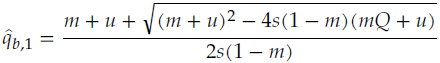

These two roots exist only if

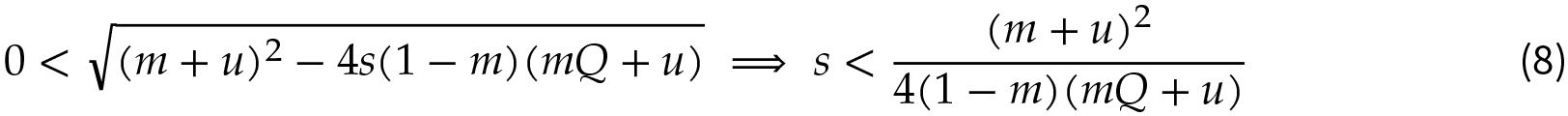

which provides us with an estimate of the upper bound on *s* for the presence of three equilibria.

The derivative of Equation 7 is 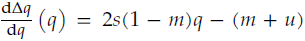, and a equilibrium will be stable if 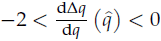. From this, it can be easily shown that 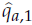 is stable and 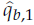 is unstable.

**Strong-selection approximation**. In order to determine the lower bound on *s* for the presence of three equilibria, we assume that selection is strong enough such that *u/s* ≈ 0 and *Q/s* ≈ 0. Therefore,

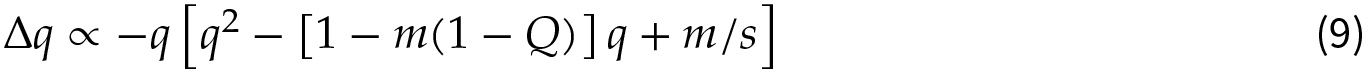

and the equilibria can be described as

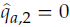

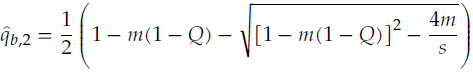

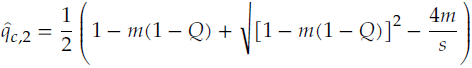

The latter two equilibria will exist only if

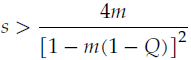

which provides us an estimate of the lower bound forthe presence of three equilibria.

The derivative of Equation 9 is 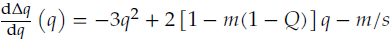, and it can be easily shown that 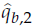 is unstable and 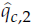 is stable.

**Validity of approximations**. By substituting 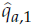 and 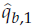 back into Equation 6, we obtain 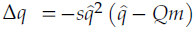. Thus, Δ*q* ≤ 0, which indicates that 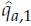 overestimates 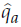 and that 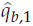 underestimates 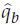. By substituting 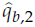 and 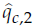 back into Equation 6, we find that 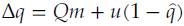. Thus Δ*q* ≥ 0, which indicates that 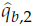 overestimates 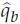 and that 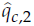 underestimates 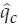. However, the error in our approximations is slight (Figure 3).

**Figure 3:**
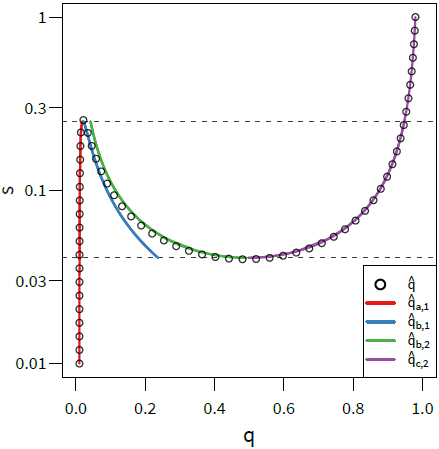
Our recessive-blindness equilibria approximations are accurate. The approximations developed
in this paper (solid lines) are a good fit for calculated values of *s* that result in equilibrium for a given *q* (circles) using Equation 3. The dashed lines are our approximate bounds for the existence of three equilibria (i.e. for small and large values of *s*, there is one equilibrium; for intermediate values of *s* there are three possible equilibria). Other parameters are *m* = 0.01, *u* = 10^−6^, and *Q* = 0.01.

**Dynamics**. Based on these approximations, the dynamics of the recessive-blindness system can be summarized as follows. First, there are three possible equilibria: 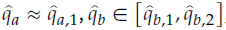, 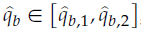 and 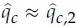. Second, there are four possible equilibria configurations: 1,2a, 2b, and 2c.

Case 1,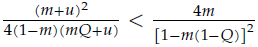: only one equilibrium exists,and it is stable. The population will always evolve towards it.

Case 2,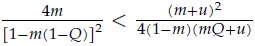: depending on the strength of *s* this case may have one of three possible configurations:

Case 2a,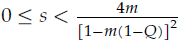: Only one equilibrium exists, 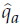 and it is stable. The population will always evolve towards it.

Case 2b, 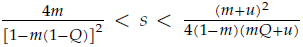: All three equilibria exist; 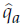 and 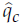 are stable, while 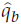 is unstable. If the population starts below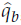, it will evolve towards 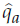. If it starts above 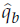, it will evolve towards 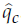.

Case 2c, 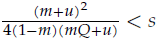 only one equilibrium, 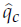, exists, and it is stable. The population will always evolve towards it.

Furthermore if *q*_0_ = *Q*, the selection-threshold for blindness to be established in the cave is

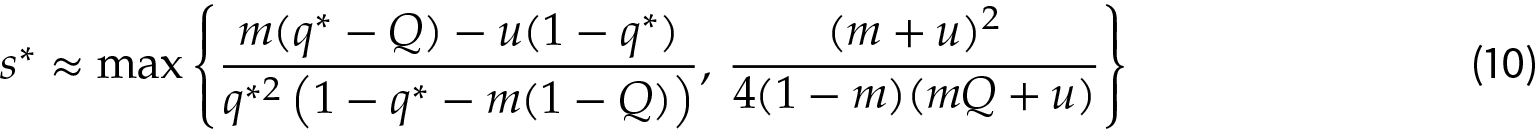

where *q*\* is the allele-frequency threshold.

### Finite-Population Simulations

**Constant migration**. To investigate the impact of drift on our recessive-blindness model, we simulated diploid populations of size *N* = 1000 by modifying our life cycle (Equation 1) to include a finite population:

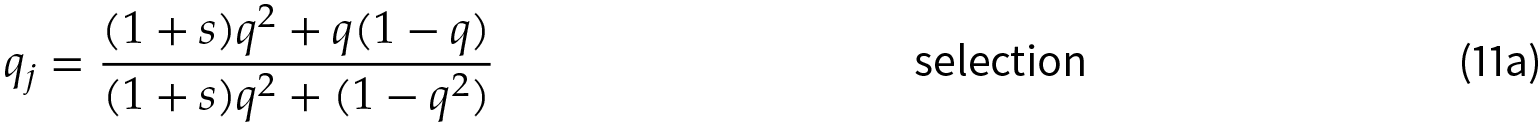

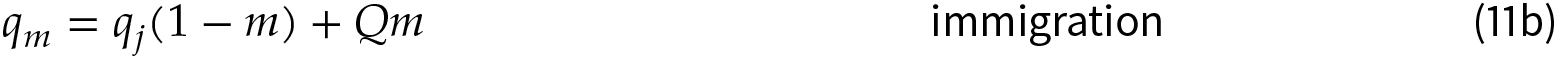

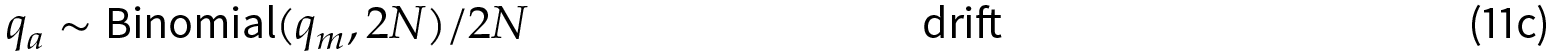

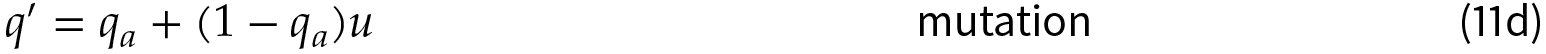

Here the adult population consists of 2*N* alleles sampled with-replacement from the post-immigration gene pool.

For every simulation, *u* = 10^-6^, *Q* = 0.01, and *q*_0_ = *Q*. We varied *s* from 10^-6^ to 10^2^ and *m* from 10^-8^ to 1. We simulated 100 replicates for each combination of parameters; simulations were conducted for 10,000 or 5,000,000 generations. For each set of parameters, we recorded the average *q′* frequency across these 100 populations at specific time points.

Our simulation results for finite populations are qualitatively similar to our analytical results for infinite populations. For high migration rates, the average allele frequency is similar to the infinite model, except that drift allows some populations that have three equilibria to evolve blindness (Figure 4B). However, at low migration rates (*m* < *u/Q* = 10^-4^), populations have low average frequency of *b* at 10,000 generations, unless *s* > 1. As immigration decreased, these populations became dependent on *de novo* mutations to produce *b*, which is a slow process. At 5 million generations, which is close to the estimated age of cavefish populations (Gross, 2012), the average allele frequency is a better match to the results from the the infinite-population model (Figure 4C); although, selection is ineffective for *s* < 1/2*N* = 5 × 10^-4^.

**Figure 4:**
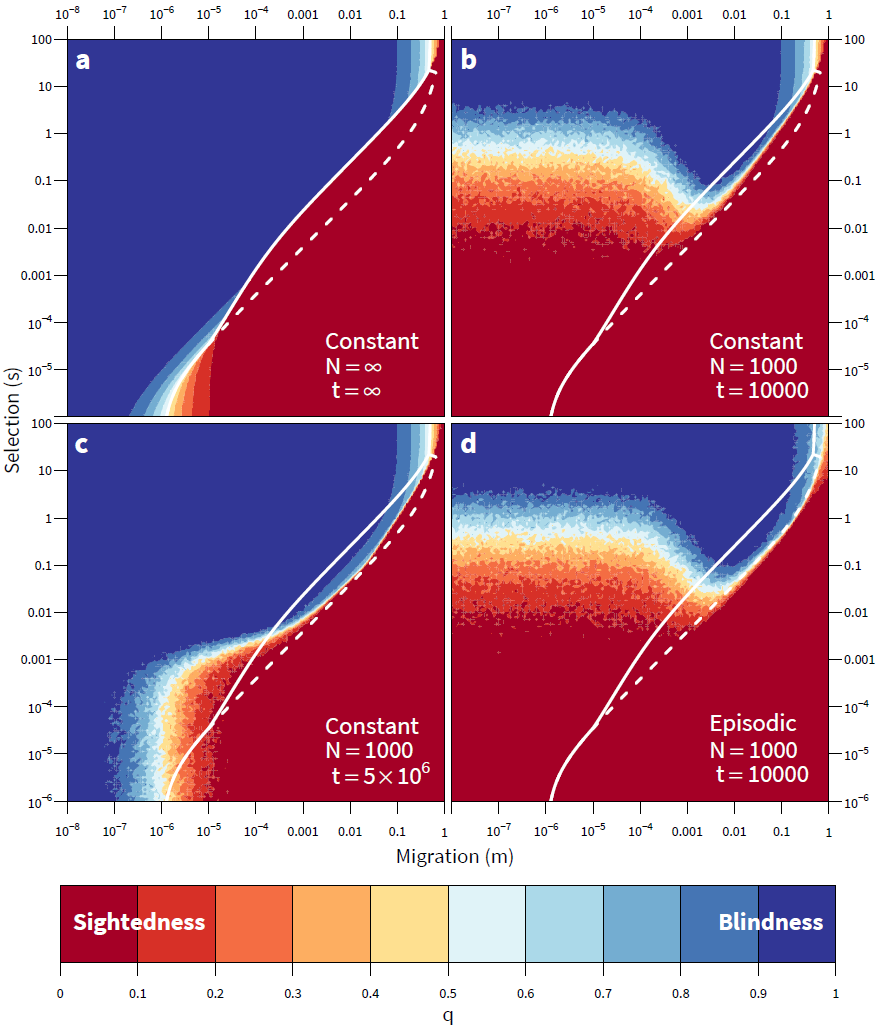
Populations evolve blindness in the face of immigration only with the help of strong selection.
(a) The equilibrium frequency of the blindness allele (*b*) for an infinite population, and (a–d) average frequencies of *b* after *t* generations in finite populations (100 replicates) with either constant or episodic migration. Colors correspond to the frequency of the blindness allele (*b*) for a given combination of selection (*s*) and migration (*m*), where blue is high frequency (blindness evolved) and red is low (blindness did not evolve). The solid white line corresponds to 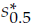 The area between the solid and dashed lines corresponds to the region where three equilibria exist. Other parameters are *u* = 10^−6^, *Q* = 0.01, and *q*_0_ = *Q*.

**Episodic migration**. Because cave and surface populations may be connected intermittently due to flooding, we simulated periods of immigration followed by periods of isolation following a first-order Markov process. The probability of switching between from isolation to immigration or vice versa was 10% in each generation. Results for the intermittently connected simulations were nearly identical to previous simulations, with the exception that at high levels of migration and selection, drift was more effective in increasing average allele frequencies (Figure 4D).

## Discussion

The evolution of blindness in caves has been hypothesized to result from relaxed selection and mutation pressure and/or positive selection for alleles that result in eye loss. However, the degree to which these factors interact and the theoretical level of selection required to induce blindness have not been quantified previously. Here we show that in case of low level immigration into a cave, blindness will eventually evolve, due to mutation and immigration of a few blindness alleles. This result fits the suggestion of some previous hypotheses: relaxed selection can result in blind populations. However, for blindness to occur in these conditions requires a significant amount of time. It is more likely that selection is much stronger than previously anticipated, allowing blindness to be produced in caves over a relatively short period of time. Furthermore, if levels of immigration are moderate to high, strong selection is necessary to produce blind populations regardless of time.

Interestingly, although cave populations are likely small, drift is only essential to the evolution of blindness in the cave population in a limited range of combinations of selection and migration for which we find three equilibria. When immigration is low, low levels of selection can lead to blindness (lower left of Figure 4A); however, in finite populations stronger selection is required to overcome the effects of drift (lower left of Figure 4C).

The amount of selection required for blindness to evolve depends on the migration rate and the level of dominance of the blindness allele (Figure 2). For example, if *Q* = 0.01 and *h* = 0, the amount of selection needs to be about 25 times the migration rate for a blind allele to become the major allele. Conversely, if *h* > 1/3, it only needs to be about 3 times. The situation is reversed when we look at fixation. If *h* = 0, selection needs to be about 100 times the migration rate for the frequency of the blind allele to exceed 99% in the cave. And if *h* = 1, it needs to be 10,000 times greater than the migration rate. If we focus on phenotypes instead, we see that dominant alleles need lower levels of positive-selection to impact the population (Figure 2).

The magnitude of selection coefficients required by our model to produce blindness given modest levels of immigration are comparable to observations in many species. Levels of selection sufficient to produce selective sweeps in wild populations range from 0.02–0.7 (Sáez et al., 2003; Schlenke and Begun, 2004; Wootton et al., 2002; Nair et al., 2003). Estimated selection coefficients for drug resistance in *Plasmodium falciparum* were 0.1–0.7, leading to fixation in 20–80 generations (Wootton et al., 2002; Nair et al., 2003). Fora major advantageous allele, the average value of *s* has been estimated as 0.11 in plants and 0.13 in animals (Rieseberg and Burke, 2001; Morjan and Rieseberg, 2004). Recent work has suggested that eye development imposes a high metabolic cost, particularly for juveniles (Moran et al.,2015). In a food-limited environment this cost could lead to strong selection, but the precise degree of this selection is unknown. The well-studied three-spine stickleback (*Gasterosteus aculeatus*) exhibits similarstrongselection in a novel environment. In experiments isolating armored sticklebacks in freshwater pools, armor was lost within a few generations due to relaxed selection for defense and positive selection for the lower cost of development in unarmored fish (Barrett et al., 2008). Estimates of selection in this species have ranged from 0.13–0.16 (Terekhanova et al., 2014).

The selection coefficient of a blindness allele is determined not only by the amount of energy saved by not having a visual system but also by any other pleiotropic effects, such as enhancement to feeding ability (Jeffery, 2005). If an allele produces multiple, adaptive phenotypes, its selection coefficient is more likely to be high enough to promote local adaptation and differentiation between cave and surface populations. Genotype-dependent dispersal (Edelaar and Bolnick, 2012; Bolnick and Otto, 2013) is one possible pleiotropic effect of blindness mutations that has not been considered in recent research on cavefish. Ninety years ago, Lankester (1925) proposed that blindness evolves in caves because fish with eyes may be attracted to sources of light and preferentially leave caves. In our model, emigration of sighted individuals would be equivalent to increasing the selection coefficient, *s*, because individuals with *B* alleles would systematically leave the cave. Even a low level of preferential emigration, e.g. 1%, would provide a significant boost to local adaptation and the evolution of blindness in caves. It is quite possible that in some species genotype-dependent dispersal combined with lower development costs promotes the elimination of sight in caves despite the immigration of sightedness alleles from the surface.

While we have drawn conclusions about a single locus, multiple genes are involved in eye development and sight. Loss-of-function mutations to any of these genes could produce blindness in caves. Linked genes would effect our model by increasing the effective mutation rate of a sighted haplotypetoa blind haplotype, reducing the amount of selection required for the evolution of blindness. Unlinked genes would provide more opportunities for drift to assist the evolution of blindness in caves.

We conclude that in most cases strong selection is necessary for the evolution of blind populations in caves. This result is consistent with two different observations of cavefish: (1) phototactic fish may leave caves, effectively selecting for the maintenance of mostly blind fish, and (2) the metabolic cost of eyes is very high. Additionally, the model and results presented in this paper are applicable beyond the evolution of cave populations, expanding existing migration-selection balance theory. We have developed approximations that allow us to understand the interaction of selection, migration, and mutation. Through simulation we have incorporated genetic drift into the model and determined that in some situations it can enhance the power of selection to drive local adaptation. Periods of isolation can also be important in these situations.

## Acknowledgments

This work was supported by Arizona State University’s School of Life Sciences and Barrett Honors College. Steven Wu, David Winter, Kael Dai, Michael Rosenberg, and Phil Hedrick provided helpful feedback on this manuscript.

## Appendix

All the proofs below were validated in Mathematica (Wolfram Research, Inc., 2015).

### Lemma 1.

*If m* > 0 *or u* > 0, 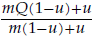 *is a possible equilibrium, and there is no equilibrium less than it. If m = u* = 0, 0 *is an equilibrium*.

*Proof*. Case 1. Let*f*(*q*) = *q*′- *q* represent the change in allele frequency over one generation (Equation 1). Let 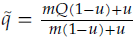.if *s* = 0 and *m* > 0(or *u* > 0), 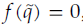, and therefore 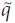 an equilibrium forthese parameters. Furthermore, if *s* ≥ 0, 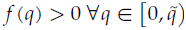. Therefore, there is no equilibrium lowerthan 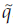.

Case 2. Let *m* = *u* = 0,*f*(0) = 0.

### Lemma 2

1 – *m*(1 – *u*)(1 – *Q*) *is a possible equilibrium, and there is no equilibrium greater than it*.

*Proof*. Let 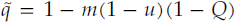 and *h* = 0. Since 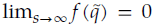,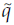 is a potential equilibrium. Furthermore, if 0 ≤ *h* ≤ 1 and *s* ≥ 0,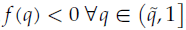. Therefore, there is no equilibrium higher than 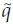.

The derivation of a tighter upper bound can be achieved by not assuming *h* = 0; however, we do not report it at this time.

### Lemma 3

*Let s* > 0. *Let m* > 0 *or u* > 0. *If h* ≥ 1/3 *or if h* < 1/3 *and* 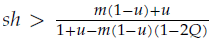, *g*(*q*) *(Equation 3) has exactly one root in* [0,1].

*Proof*. Let *m* > 0 or *u* > 0. Then *g*(1) < *g*(0) and *g*(1) ≤ 0 ≤ *g*(0). By the intermediate value theorem there is at least one root in [0,1]. Let *s* > 0 and we will show that there is exactly one root for several cases.

Case 1. Let 1/2 < *h* ≤ 1. Then *g*(−∞) < 0 and *g*(∞) > 0. By the intermediate value theorem, *g*(0) has at least one root below 0, between 0 and 1, and above 1. Since *g*(0) is cubic, it can have at most 3 roots; therefore, there is exactly one root in [0,1].

Case 2. Let *h* =1/2. *g*(*q*) reduces to a quadratic equation with one root less than 0 and exactly one root in [0,1].

Case 3. Let 1/3 ≤ *h* ≤ 1/2. 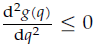, and *g*(*q*) is concave in [0,1]. Thus *g*(*q*) has exactly one root in [0,1].

Case 4. Let 0 ≤ *h* ≤ 1/3 and 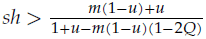.then 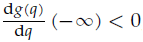,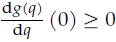,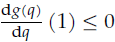, and 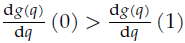. By the intermediate value theorem, theremust be a localminimumin (-∞, 0]and and a localmaximumin [0, 1]. Thus *g*(*q*) has exactly one root in [0, 1].

**Derivation of Equation 5.** In order to derive Equation 5 we first assume that *u* =0. Then

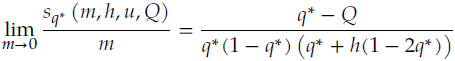

However, we also need to determine when Δ*q* has only one root. First we approximate Δ*q* by a second-order Taylor series near *q* = 0.

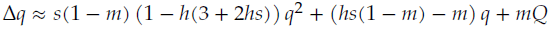

Next we find

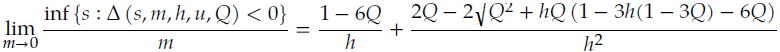

where Δ (*s,m, h, u, Q*) is the discriminant of the Taylor approximation.

Equation 5 is the maximum of these two values.

